# RNA-Dependent Synthesis of Mammalian mRNA: Identification of Chimeric Intermediate and Putative End Product

**DOI:** 10.1101/071266

**Authors:** Sophia Rits, Bjorn R. Olsen, Vladimir Volloch

## Abstract

Our initial understanding of the flow of protein-encoding genetic information, DNA to RNA to protein, a process defined as the “central dogma of molecular biology”, was eventually amended to account for the information back-flow from RNA to DNA (reverse transcription), and for its “side-flow” from RNA to RNA (RNA-dependent RNA synthesis, RdRs). These processes, both potentially leading to protein production, were described only in viral systems, and although putative RNA-dependent RNA polymerase (RdRp) was shown to be present, and RdRs to occur, in most, if not all, mammalian cells, its function was presumed to be restricted to regulatory. Here we report the occurrence of protein-encoding RNA to RNA information transfer in mammalian cells. We describe below the detection, by next generation sequencing (NGS), of a chimeric doublestranded/pinhead intermediate containing both sense and antisense globin RNA strands covalently joined in a predicted and uniquely defined manner, whose cleavage at the pinhead would result in the generation of an endproduct containing the intact coding region of the original mRNA. We also describe the identification of the putative end product of RNA-dependent globin mRNA amplification. It is heavily modified, uniformly truncated at both untranslated regions (UTRs), terminates with the OH group at the 5’ end, consistent with a cleavage-generated 5’ terminus, and its massive cellular amount is unprecedented for a conventional mRNA transcription product. It also translates in a cell-free system into polypeptides indistinguishable from the translation product of conventional globin mRNA. The physiological significance of the mammalian mRNA amplification, which might operate during terminal differentiation and in the production of highly abundant rapidly generated proteins such as some collagens or other components of extracellular matrix, with every genome-originated mRNA molecule acting as a potential template, as well as possible implications, including physiologically occurring intracellular PCR process, iPCR, are discussed in the paper.

## INTRODUCTION

Previously, a mechanism was proposed for RNA-dependent amplification of mRNA in mammalian cells (Fig. 1, “model” panel). In this postulated process, mature mRNA is transcribed by RdRp. Such transcription initiates within the 3’ poly(A) region of mRNA and produces an antisense strand containing poly(U) at the 5’ end and terminating at the 3’ end with a complement of the 5’ terminus of the mRNA molecule. The subsequent copying of the antisense strand was postulated to occur via self-priming and extension of its 3’ terminus. Cleavage within a single-stranded loop of the resulting pinhead molecule would then produce sense strand terminating in poly(A) and a 3’-truncated antisense strand. If self-priming occurs within the segment corresponding to the 5’untranslated region (UTR) of mRNA, the resulting sense strand would contain the entire protein coding information of the original mRNA and would translate into a polypeptide indistinguishable from the conventionally produced protein. If self-priming occurs within the segment corresponding to the coding region of mRNA, the translational outcome would depend on the location of the first eligible AUG initiation codon; if it is “in frame” with the initial coding sequence, a C-terminal region of the original protein would be produced (Volloch, 1996; 1997).

**Figure 1.**
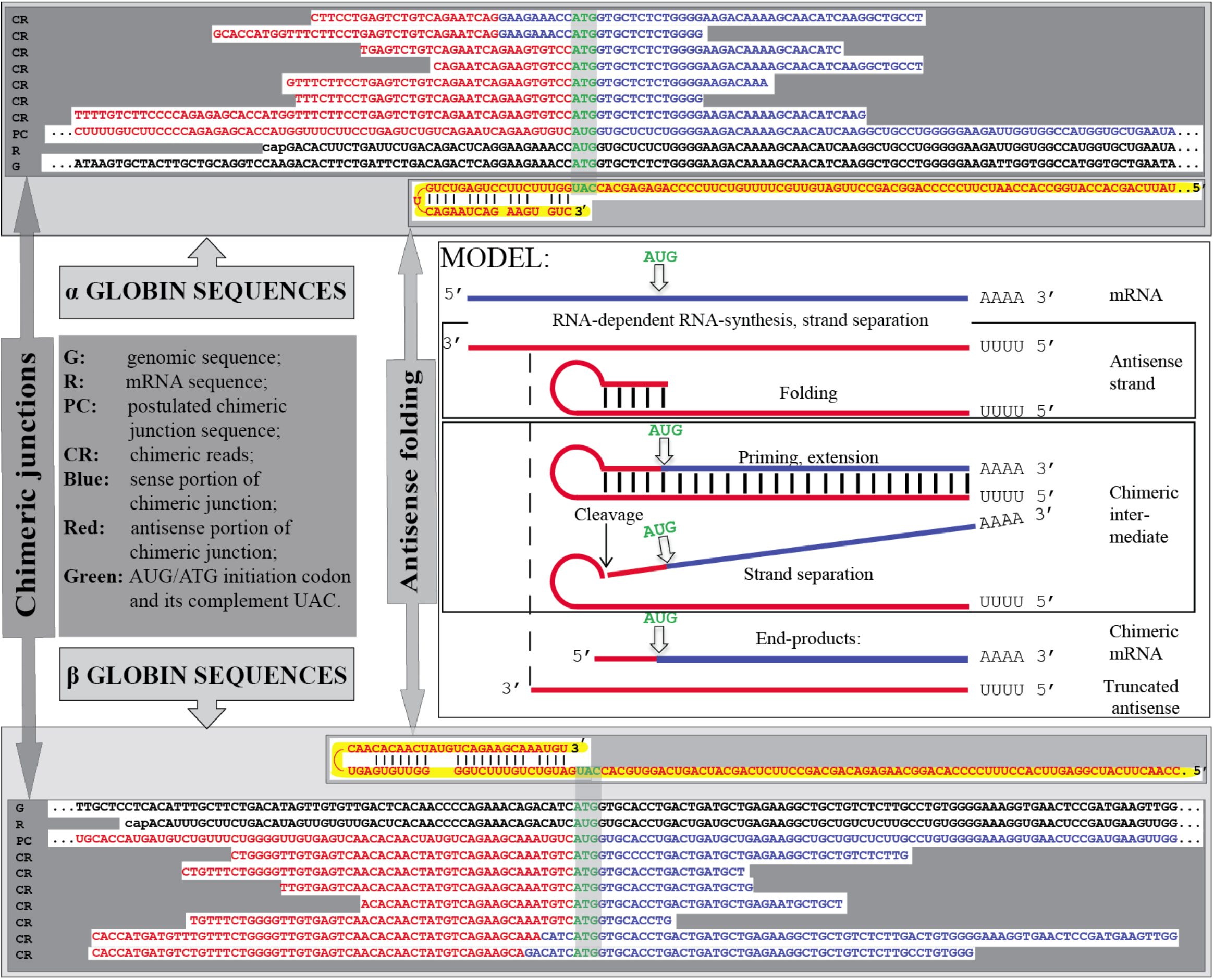
Detection of chimeric junctions containing both sense and antisense globin sequences in RNA from anemic spleen.

The proposed mechanism described above originated in studies of globin mRNA expression in differentiating murine erythroid cells (Volloch et al., 1996). In these studies antisense strands were detected and characterized by sequencing following ligation-mediated RNA amplification. It was shown (Volloch et al., 1994) that a segment of antisense strand corresponding to the 5’UTR of globin mRNA contains two complementary elements, one of them 3’terminal (Fig. 1, “folded antisense RNA” panels; Fig. 2). Model experiments demonstrated that this complementarity, although imperfect in that it contains mismatches and utilizes G/U pairing, enables self-priming and extension of the antisense strand and that this process crucially depends on the strict 3’terminal localization of one of the complementary elements (Volloch et al., 1994). Studies also showed (Volloch et al., 1994) that the self priming-enabling complementarity within the globin antisense strand is preserved throughout a large evolutionary distance with conservation of not only the occurrence of complementary elements but also their positions within the segment corresponding to the 5’UTR of mRNA. Although nucleotide sequences in the 5’ UTRs of adult globin mRNAs diverged substantially during evolution, the complementary relationship of the 3’ terminal and the internal elements of an antisense strand remained preserved.

**Figure 2.**
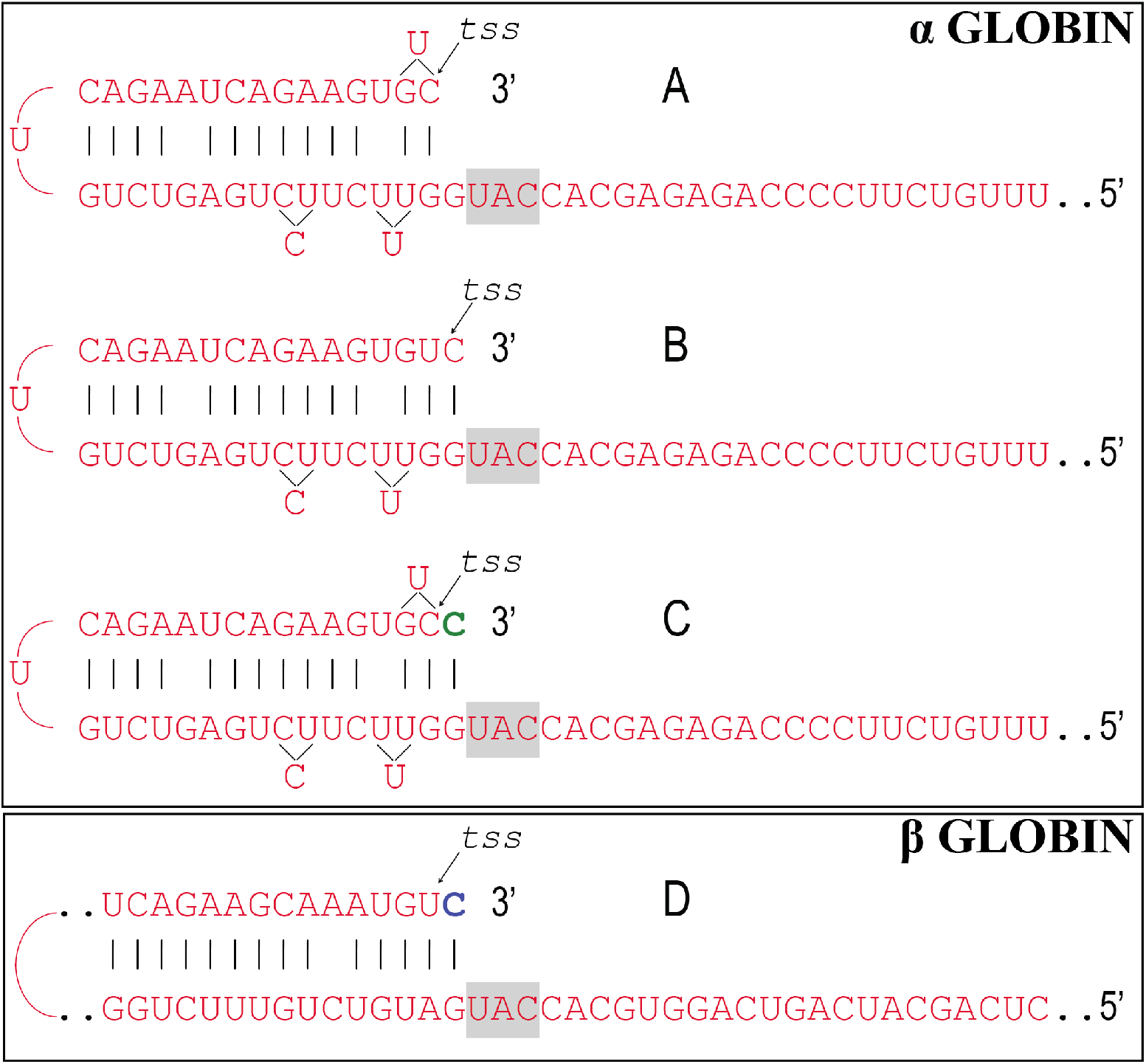
RNA-dependent RNA polymerase can transcribe the cap”G” of mRNA. Folding of: antisense α globin (A-C) or β globin (D) RNA. *tss*: nucleotide corresponding to the transcription start site of mRNA. Green: transcript of the cap”G”. Blue: transcript either of cap”G” of mRNA or of complementary “G” during the 3’ extension of folded antisense RNA. Highlighted: complement of the AUG initiation codon.

This proposed mechanism remained, until now, largely hypothetical because no convincing evidence in support of the mechanism could be obtained. Indeed, the detection and characterization of globin antisense RNA (Volloch et al., 1996) does not constitute decisive evidence for the occurrence of the proposed RNA amplification mechanism since it could have been generated for some other purposes, and the identification of its predominant 3’-terminal truncation in a position consistent with the postulated cleavage at the 3’end of a singlestranded loop of the pinhead molecule that would define the 5’ terminus of the end product (Volloch et al., 1996) is, on its own, an indirect indication. But what would constitute convincing evidence? At the core of the proposed mechanism is the well-defined extension of a self-primed antisense strand. Therefore, the detection of its major recognizable attribute, an antisense molecule extended into a sense strand in a predicted and uniquely defined manner, would provide decisive support for the proposed mechanism for the mRNA amplification process. Two types of postulated RNA molecules would, due to their intrinsic structure, contain such evidential information. One is the double-stranded/pinhead chimeric intermediate consisting of a sense and an antisense strand covalently joined in a precise and defined way, a direct precursor of the end-product. The other is a chimeric end-product containing an antisense fragment at its 5’terminus. In both cases the region of interest is the junction between the sense and the antisense portions. Such junctions of a chimeric intermediate would be readily identifiable if sufficiently long sequences could be obtained. Even the postulated chimeric end-product would be clearly and predictably distinguishable in its junction sequence from the corresponding sequences of conventional mRNA of genomic origin due to the imperfect complementarity of the self-priming elements of an antisense strand (Volloch et al., 1994; 1996). Therefore, in the present study we tested, using newly available technologies, for the occurrence of such chimeric sense/antisense junction sequences in murine erythroid cells.

## RESULTS

### Substantiation of the proposed mechanism: detection of chimeric junction sequences

The occurrence of chimeric sense/antisense globin RNA molecules was tested by next generation RNA sequencing. Cytoplasmic RNA from spleen cells of anemic mice was used to generate two types of sequencing libraries, Illumina TruSeq RNA library and NEB Ultra Directional RNA library for Illumina, that were analyzed on a MiSeq sequencer. The resulting reads were aligned with or blasted against appropriate references, and sequences of the fragments of interest were extracted from raw data and analyzed. A selection of chimeric fragments detected for both alpha and beta globin RNA is presented in Fig. 1, together with alpha and beta globin genomic reference sequences, conventional globin mRNA sequences, postulated chimeric sequences, and folded antisense sequences whose 3’ extension would generate predicted chimeric sequences. The figure also contains a schematic summary of the postulated RNA amplification process that results in chimeric molecules. As shown in Fig. 1, two types of chimeric reads were obtained: those originated from “full-size” antisense RNA and those (two upper reads for alpha globin and two bottom reads for beta globin) consistent with originating from a 3’-truncated or prematurely terminated antisense RNA retaining a portion of its terminal complementary element apparently sufficient for self-priming.

The key concern about the observed chimeric junction sequences is their authenticity, i.e. whether they are produced by the proposed mechanism or are an artifact. There is only one obvious process by which the observed sequences could have arisen artificially. It was described in an earlier study (Volloch et al., 1994) where it was shown that after the completion of the synthesis of globin antisense cDNA strand by reverse transcriptase, it can, using the very same complementary elements described above, self-prime the synthesis of a sense strand. This process was absolutely dependent on the presence in cDNA of two complementary elements and on RNase H activity for the removal of template RNA. Several lines of reasoning strongly indicate that the observed chimeric RNA fragments are not the result of such a process. First, reverse transcriptases employed in preparation of sequencing libraries were RNase H-negative mutants. Second, even if such chimeric structures would have been generated by reverse transcription, they would have only one “open” end and the procedure of library preparation, which requires adaptor ligation to both ends, would select against them. Third, chimeric sequences were observed not only with Illumina True Seq RNA library but also with NEB Ultra Directional RNA sequencing library, where the second cDNA strand is eliminated as a part of the library construction. Fourth, and the decisive argument for authenticity, is the intrinsic geometry of the observed chimeric reads: all obtained chimeric sequences shown in Fig. 1 have, when folded by matching the complementary elements discussed above, 3’-protruding globin-specific segments, a feature incompatible with its possible artifactual origin. Fifth, moreover, some of the chimeric fragments sequenced, in addition to exhibiting 3’ globin-specific protrusions, also lack the entire internal complementary element (for example, middle reads for both alpha and beta globins in Fig. 1), another impossibility within the framework of the artifactual cDNA generation process.

All chimeric junction sequences observed corresponded to those postulated on the basis of the proposed mRNA amplification model with the unexpected exception that, compared with the sequence prediction based on the folding of the alpha globin antisense strand depicted in Fig. 1, all alpha globin chimeric sequences obtained contained one extra “C” at the extreme 3’ end of the antisense portion, immediately preceding the “ATG” initiation codon (Fig. 1). Its presence is trivially explained for 3’ truncated antisense RNA-originated chimeras where it is inserted during the extension phase. The explanation is more complex for “full-size” antisense RNA-originated chimeric molecules. It posits that the full-size antisense strand contains, in addition to the genome-encoded sequence and as an integral part of its primary structure, an added 3’-terminal “C” and, moreover, that this “C” is the transcript of the cap “G” of the sense strand. This conclusion is arrived at by the exclusion of possible alternatives. If the 3’ terminus of the antisense strand faithfully reflects the transcription start site (TSS) of the sense strand, i.e. does not contain an extra “C”, the only explanation for the additional “C” in the chimeric sequence is that folding of the antisense strand occurs as shown in Fig. 2A and therefore the first template nucleotide during the extension is “C”; however, with the available folding alternative depicted in Fig. 2B, this explanation is not a viable one for thermodynamic reasons. In contrast, with an additional 3’-terminal “C” as the integral part of the primary sequence, the antisense strand folding would occur as shown in Fig. 2C and account for the additional “C” in the sequence of the chimeric junction. Since the genomic sequence upstream of the TSS for alpha globin (Fig. 1) cannot account for the additional 3’-terminal “C” in the antisense strand, the only remaining possibility is that the “C” in question is a transcript of the cap “G” of the sense strand. RNAdependent DNA polymerases are known to transcribe the cap “G” despite its parallel rather than opposite (in relation to an antisense) orientation (Hirzmann et al., 1993; Volloch et al., 1995; Mules et al., 1998; Bibillo and Eickbush, 2004) and such an ability appears to be an attribute also of a mammalian RNA-dependent RNA polymerase. In the case of beta globin, the antisense folding is such (Fig. 2D) as to either accommodate the cap-transcribed “C” or, if the cap is not transcribed, to account for the “AUG”-preceding “C” generated during the extension. To distinguish between these two possibilities is not possible.

The variable length of antisense portions of the chimeric sequences obtained, some extending well into the region corresponding to the coding portion of the sense strand, as well as the amount of detected chimeric fragments, only about 0.001% of total reads, indicate that detected fragments are likely those of doublestranded/pinhead chimeric intermediates and not of the single-stranded end product. The conspicuous absence of potentially highly ubiquitous end product of amplification suggested a possibility that it might be not detectable by a reverse transcription-based method of analysis employed in the present study. In the course of this study we did, in fact, encounter a novel, highly ubiquitous and heavily modified RNA species described below that exhibited not only the above attribute of non-detectability by conventional sequencing methods but also several features consistent with its putative identity as the end product of globin mRNA amplification. It can be argued (see Discussion) that nucleotide modifications, which apparently interfere with reverse transcription of this RNA species, are introduced as the integral part of its generation process and therefore a mature amplified molecule, the end product, is never present in an unmodified conventionally detectable form.

### Identification of the putative end product of globin mRNA amplification

Deficiency of red blood cells in mammals, due to blood loss or disease, stimulates conversion of the spleen into an erythropoetic organ (Conkie et al., 1975; Bastos et al., 1977; Cao and Galanello, 2010). This splenic reaction is particularly dramatic in rodents; seven days post-induction of hemolytic anemia in mice the spleen mass increased nearly 20 fold, with basophilic erythroblasts constituting the majority of cells. Electrophoresis of total cytoplasmic RNA from anemic mouse spleen on agarose/methylmercury gels showed the presence of an increasingly pronounced band of about 550-600 nucleotides (Fig. 3; Fig 4, left panel). It was resistant to DNase and disappeared following RNase or alkaline treatments. Its emergence is associated with hemoglobin accumulation (Wojda et al., 2002): it is not present in spleen cells prior to the re-activation of erythropoiesis in this organ and increases in relative amount as more cells commit to differentiation and accumulate hemoglobin (Fig. 3). Based on the size of this RNA, its kinetics and abundance, and in light of results described below, it will be referred to as pepRNA (*putative end product* of globin mRNA amplification). Eventually, erythroid cells mature and are translocated into the bloodstream where they are present as reticulocytes constituting the majority of the cell population (Bastos et al., 1977) and where the levels of hemoglobin are maintained at a plateau (Wojda et al., 2002). During this erythroblast/reticulocyte transition pepRNA levels drop sharply, declining more than six-fold as a proportion of ribosomal RNA (Fig. 4, left panel). The pepRNA band produced no signal upon stringent hybridization of Northern blots with globin-specific probe (Volloch et al., 1996). Since nucleotide modifications can interfere with hybridization properties (Hundley and Bass, 2010), we analyzed (Crain, 1990; Pomerantz and McCloskey, 1990) the nucleoside composition of pepRNA (Fig. 4, right panel). In addition to the nucleoside peaks seen in digests of control oligo(dT)-selected RNA eluted from the same gel as pepRNA, two new peaks, marked X and Y, were evident in the profile of pepRNA. Mass-spectrometry produced mass numbers of 304 and 320 for the new peaks and standard numbers for the four conventional peaks (267, 283, 244 and 243 for A, G, U and C, respectively). Together, the new peaks comprised 18%, nearly one fifth, of all nucleoside residues.

**Figure 3.**
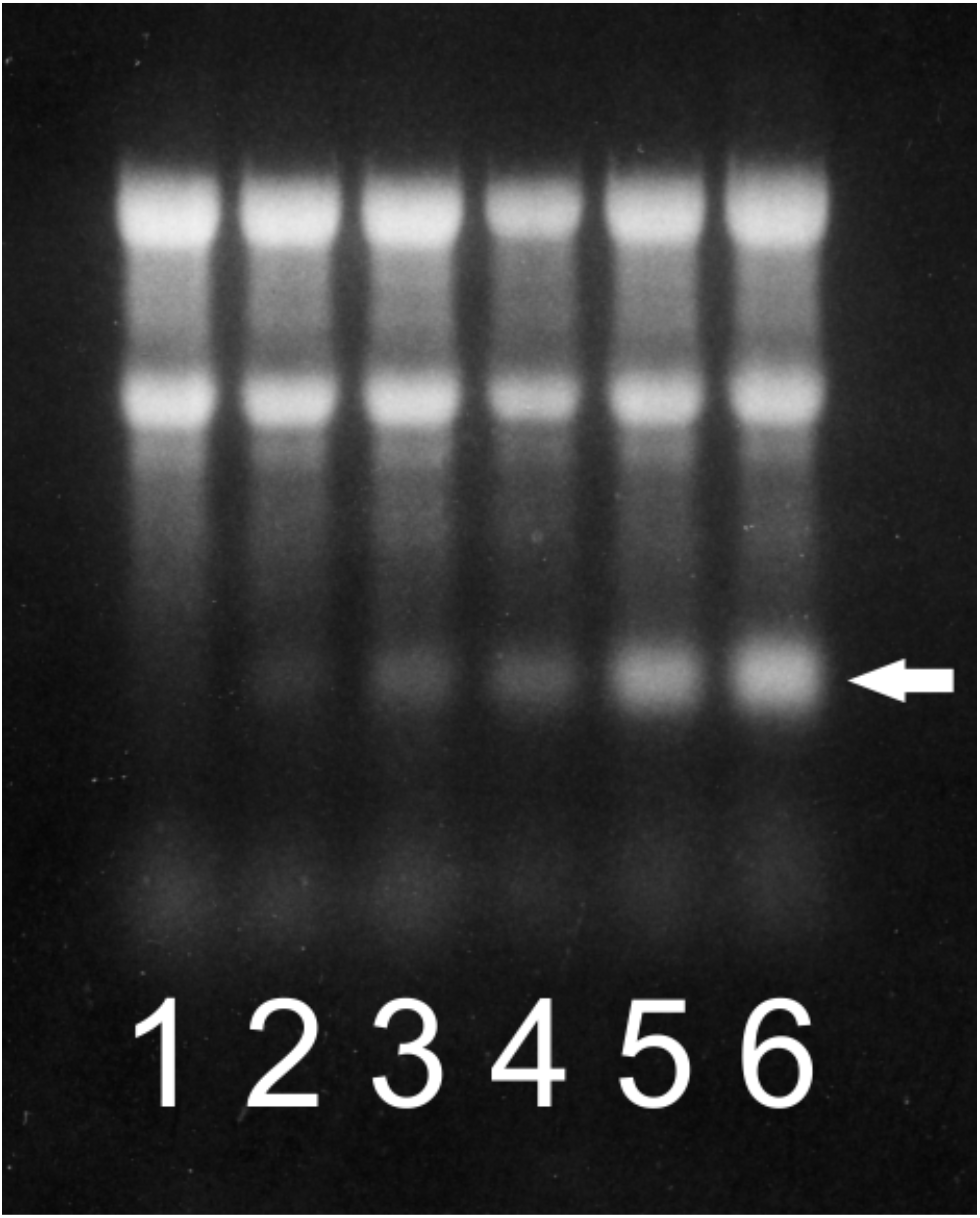
Proportion of pepRNA increases as a function of time. Ethidium bromide-stained total cytoplasmic RNA resolved on a denaturing agarose/methylmercury gel. 1-6: number of daily phenylhydrazine injections. Spleens were collected and processed 24 hours after the final injection. Two prominent upper bands in each lane: ribosomal RNA; arrow: pepRNA.

**Figure 4.**
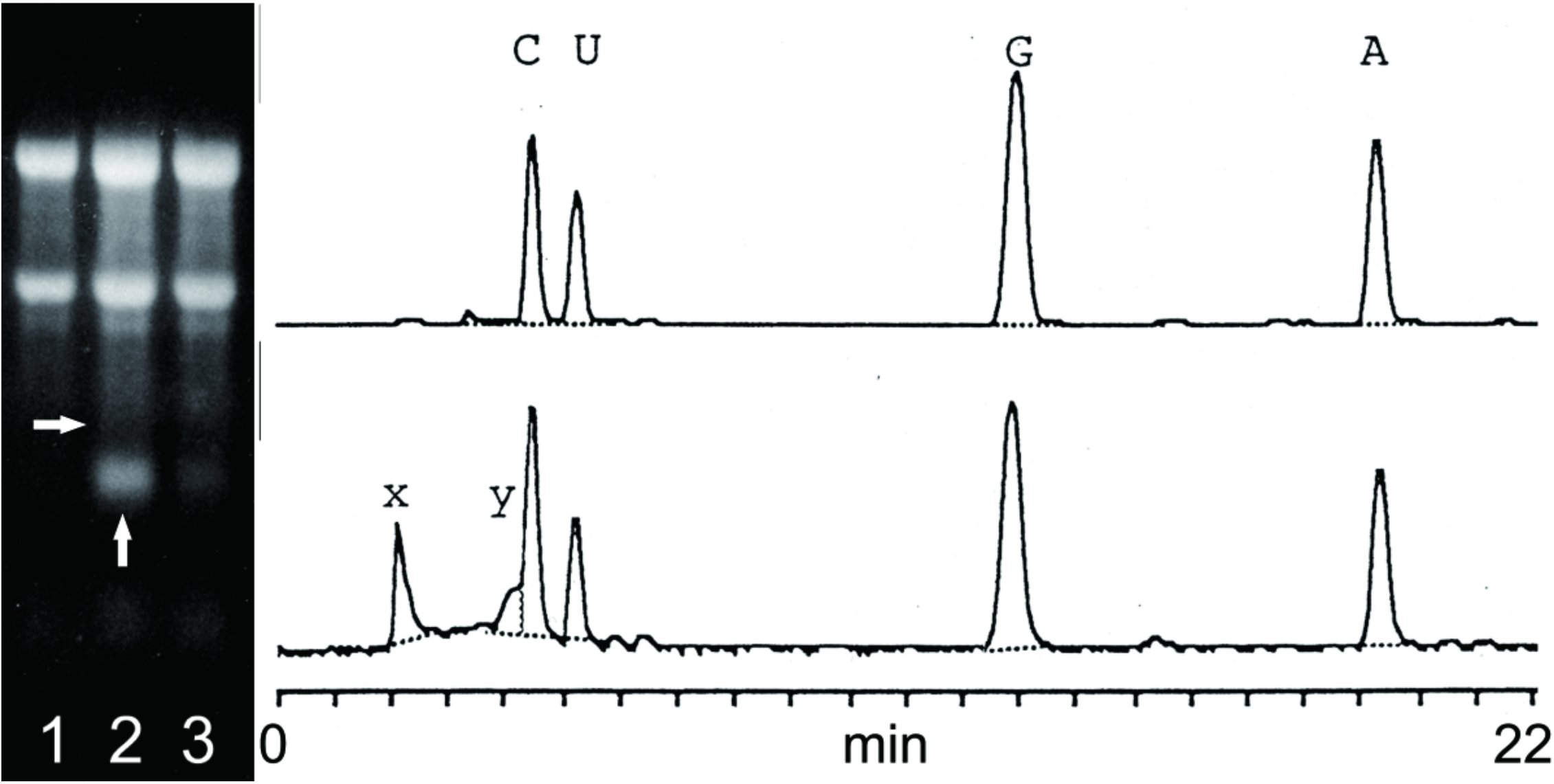
pepRNA contains modified nucleotides. Left panel: ethidium bromide-stained total cytoplasmic RNA resolved on a denaturing agarose/methylmercury gel; lane 1 – RNA from normal spleen, lane 2 –RNA from anemic spleen, lane 3 – RNA from reticulocytes of anemic mice. Two prominent upper bands in each lane: ribosomal RNA. Vertical arrow: pepRNA; horizontal arrow: estimated position of conventional globin RNA. Right panels: OD260 elution profiles of nucleoside digests of control RNA (upper panel) and pepRNA (lower panel). A, C, G, U: conventional nucleotides; X, Y: modified nucleotides.

When Northern blots were hybridized with globin-specific probes under low stringency conditions, the pepRNA band produced a signal, however, increased background precluded any claim of specificity. Therefore, we analyzed the pepRNA using hybridization with globin-specific oligonucleotide probes at relatively low stringency with added levels of specificity, such as direct nucleotide sequencing by primer extension or RNase H mapping analysis. Several synthetic oligonucleotides complementary to alpha or beta globin RNA were used as primers in reverse transcription sequencing of pepRNA and conventional globin RNA. PepRNA was eluted from gels and depleted of regular globin RNA by high stringency hybridization with immobilized globin-specific probes (Volloch et al., 1990; Volloch et al., 1991). Conventional globin RNA, isolated from reticulocytes or eluted from the same gel as pepRNA, was also purified by high stringency hybridization. Sequences obtained with all globin-specific oligonucleotide primers used and conventional globin RNA were of expected length, but all extension reactions with pepRNA, when they occurred, terminated after incorporation of only few nucleotides; when set of random primers was used with pepRNA, only short labeled fragments were produced. Sequences obtained with pepRNA and defined primers appeared to be globin, but specificity cannot be claimed due to their insufficient length.

For RNase H mapping, RNA was labeled at 5’ or 3’ ends, prehybridized with an excess of different oligodeoxynucleotide probes and treated with RNase H. Maps of fragments, obtained using the same sets of alpha and beta globin-specific oligonucleotide probes with conventional globin RNA or with pepRNA, are shown in Fig. 5. In experiments with 3’-labeled conventional globin mRNA (Fig. 5A-D), the noticeable feature was a diffuse poly(A) tail of variable length, clearly seen with the short fragments produced by RNase H cleavage. In parallel experiments with pepRNA, fragments produced by RNase H cleavage had the same electrophoretic mobility as corresponding fragments of regular globin RNA but with uniformly short poly(A) tail. In RNase H cleavage maps from experiments with regular globin RNA or with pepRNA labeled at the 5’ terminus, the pepRNA-originated fragments exhibited patterns of electrophoretic mobility on gels similar to those of corresponding conventional globin RNA-originated fragments (Fig. 5E-H). However, fragments of pepRNA were uniformly shorter than corresponding fragments obtained with regular globin RNA; apparently, 20-25 nucleotides shorter for alpha- and 30-35 nucleotides shorter for beta globin RNA probes. Oligonucleotide probes differed in their efficiency of mediating cleavage of pepRNA. When two, one alpha globin-specific and another beta globin-specific, “efficient” probes were used together, complete cleavage of pepRNA by RNase H was observed (Fig. 5I).

**Figure 5.**
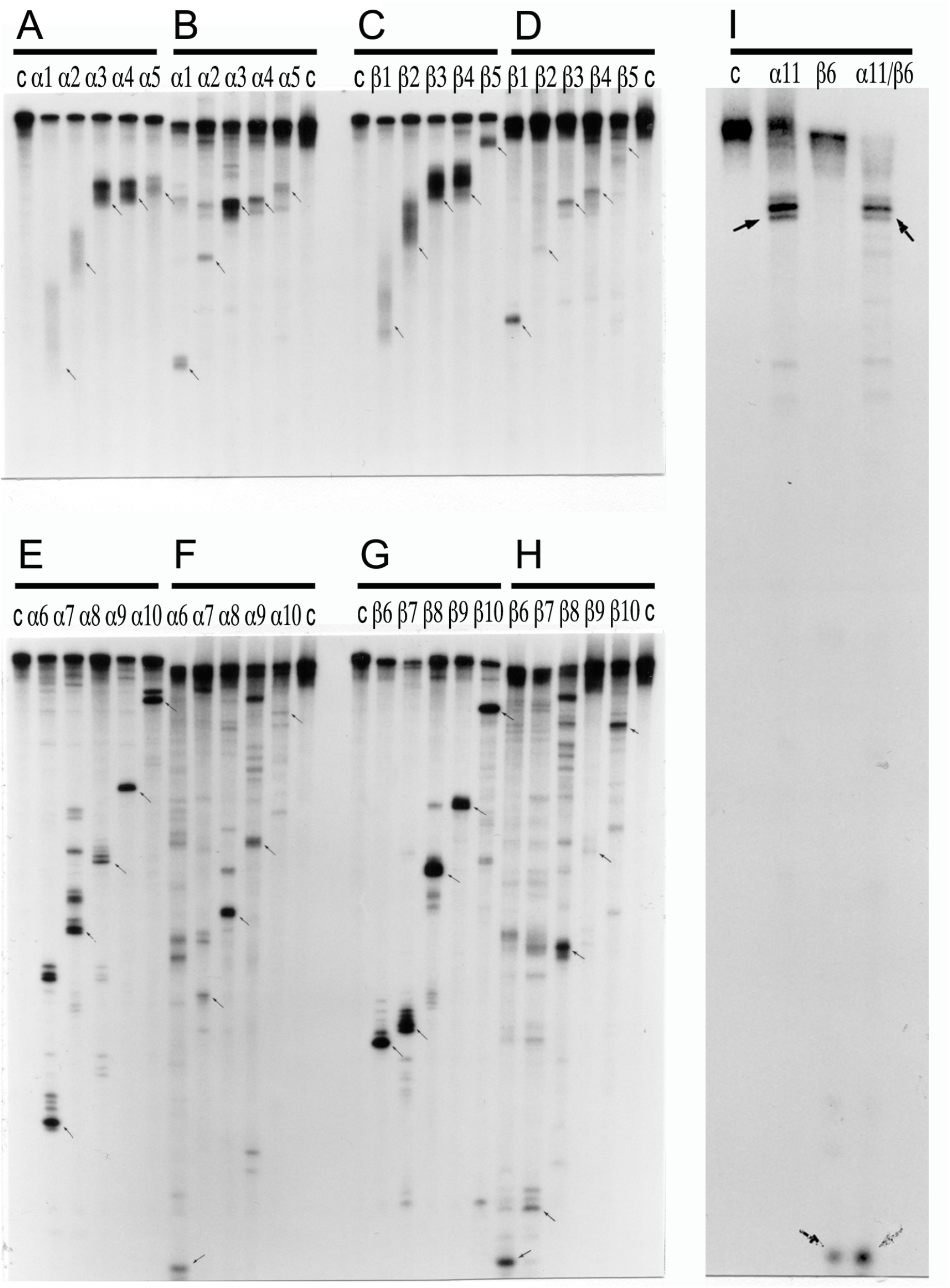
Comparative RNase H mapping of pepRNA and conventional globin mRNA. Panels A, C, E, and G: regular globin mRNA. Panels B, D, F, H, and I: pepRNA. A-D: 3’ labeled RNA. E-I: 5’ labeled RNA. α1-α11and β1-β10: alpha and beta globinspecific oligo-nucleotides used in RNase H digest assays. α11/β6: oligonucleotides used jointly in RNase H digest reaction. C: control, no oligonucleotide added.

Observations made during RNaseH mapping analysis led us to conclude that pepRNA terminates with the (OH) group at both 3’ and 5’ termini. In these experiments, the 5’ ends of the regular globin RNA and of pepRNA were labeled by polynucleotide kinase-mediated addition of radioactive phosphate. Before labeling, conventional globin RNA was pretreated with pyrophosphatase to remove the cap and with phosphatase to remove the remaining 5’phosphate and expose the (OH) group. When these pretreatments were omitted, incorporation of radioactive phosphate was suppressed apparently for the lack of substrate. No pretreatment, however, was needed for labeling of the pepRNA and the kinetics of kinase reactions were similar with pretreated regular globin RNA and non-pretreated pepRNA; when pepRNA was pretreated, there were neither increase in incorporation nor changes in the mapping outcomes. These observations indicate that the bulk of pepRNA terminates at the 5’end with the (OH) group.

3’ end-labeling of RNAs was carried out using labeled pCp and T4 RNA ligase. Similar levels of incorporation were obtained with both conventional globin RNA and pepRNA. However, the kinetics of incorporation was much slower with the latter than with the former. These observations suggest that RNA ligase may have difficulties recognizing the 3’ terminus of pepRNA as a substrate.

The above results indicate that pepRNA is a variant of globin mRNA. According to RNase H mapping, it contains a complete coding region. Whether it can be translated into globin polypeptides was tested in cell-free translation followed by gel analysis. With pepRNA, a single major translation product that could be immunoprecipitated with sheep antisera against murine adult globins and co-migrated with the translation product of conventional globin RNA was seen (Fig. 6). The possibility that the translation product of pepRNA was due to contamination with regular globin RNA was ruled out by depleting pepRNA of regular globin RNA, prior to the translation assay, by high stringency hybridization with immobilized globin-specific probes, and testing it, after binding to nitrocellulose, by high stringency hybridization with labeled globin-specific probes; this yielded a negative result.

**Figure 6.**
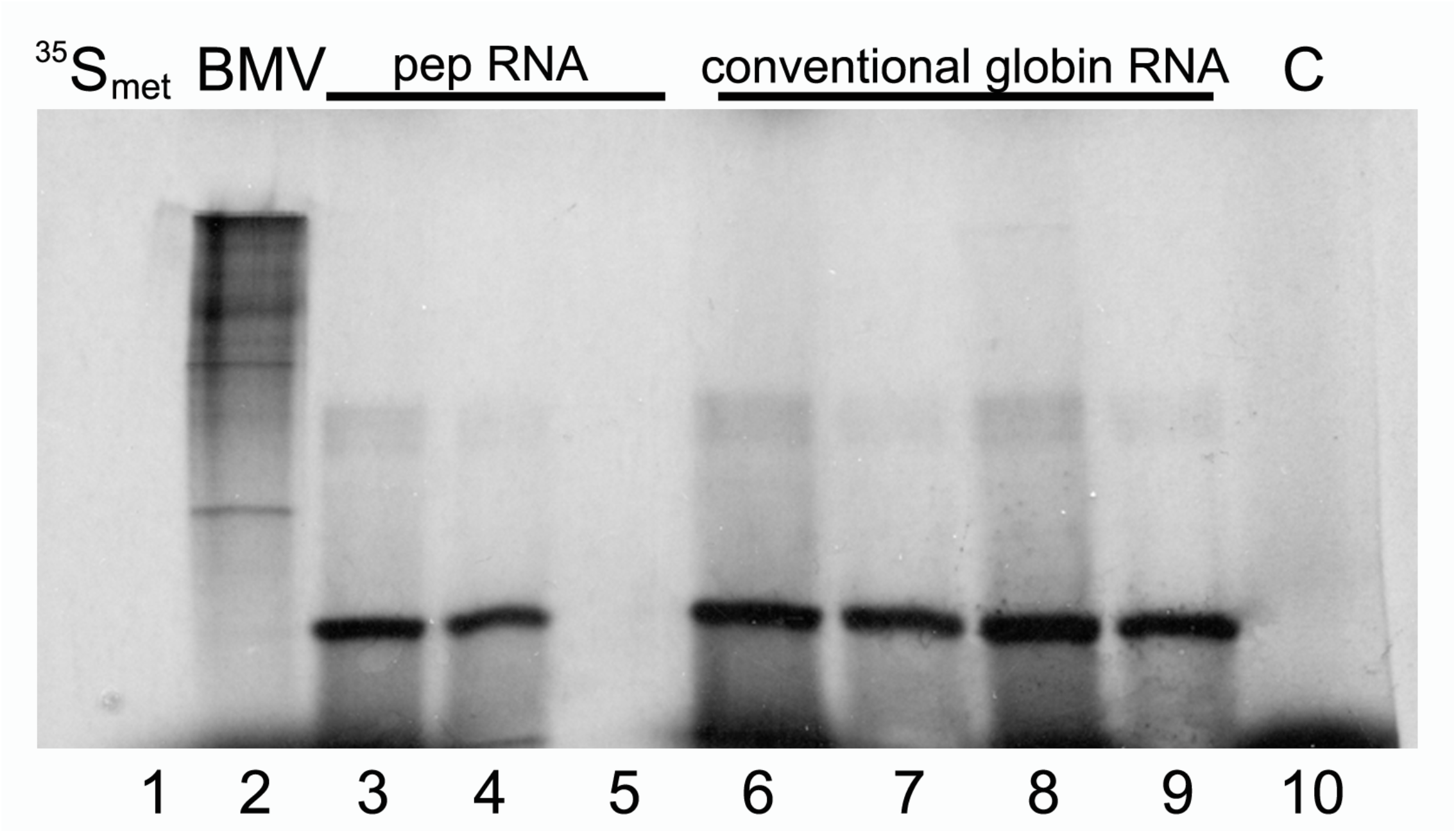
pepRNA directs synthesis of globin polypeptides in cell-free translation system. Lane 1: 35Smet, 35S-labeled methionine; Lane 2: BMV, brome mosaic virus RNA. Lanes 3-5: pepRNA; lanes 6, 7: reticulocyte globin RNA; lanes 8, 9: conventional globin RNA from anemic spleen. Lanes 4, 7, 9: translation product immunoprecipitated with sheep antisera against adult mouse globins; lane 5: preimmune serum was used in immunoprecipitation. C: control, no RNA added.

## DISCUSSION

The results described above strongly support the proposed mechanism for RNA-dependent amplification of globin mRNA. The decisive substantiation of the mechanism comes from identification of its major recognizable attribute, chimeric RNA consisting of both sense and antisense sequences covalently joined in a predicted and uniquely defined manner. These results constitute conclusive evidence for the occurrence of RNA-dependent synthesis of mRNA in mammalian cells. The scope of this process is indicated by the vast cellular amount of the putative end product, at its peak constituting 15% of rRNA, unprecedented for a conventional genome-originated mRNA. All results obtained, chief among them the capacity to direct synthesis of globin polypeptides, are consistent with the predictions of the proposed mechanism and, cumulatively, indicate that pepRNA *is* the end product of RNA-dependent synthesis of globin mRNA. The term “putative” is used here only in deference to its inability to hybridize with a probe and to support reverse transcription and, consequently, sequencing. These properties, apparently conferred by nucleotide modifications, have two possible explanations. One is that modifications interfere with the formation of complementary bonding. Another is that modifications make this type of RNA “sticky” and mediate the formation of hard to unfold complexes. The second explanation is supported by the following observations. When cytoplasmic RNA from anemic spleen cells is resolved on agarose/formaldehyde gel, transferred to a membrane and hybridized with globin-specific labeled RNA probe, in addition to an expected signal with conventional globin mRNA, an order of magnitude stronger signal is seen with ribosomal bands. There is no signal with similarly processed RNA from normal spleen. It appears that pepRNA form complexes with rRNA where “sticky” modifying groups are sequestered enabling hybridization with RNA probe, an observation that explains translation of pepRNA despite its inability to hybridize in free form: engagement of a codon with rRNA at the ribosome A site apparently makes its presentation to and hybridization with an anticodon feasible. pepRNA/rRNA interaction appears to be non-specific. When anemic spleen is extracted with guanidinium (Chomczynsky and Sacchi, 1987; 2006), pepRNA is largely lost from the RNA fraction, yet when phenol/chloroform-extracted cytoplasmic RNA from anemic spleen is re-extracted with guanidinium, pepRNA is fully retained. It appears therefore that upon disruption of tissue, pepRNA complexes with DNA, the predominant nucleic acid species amenable, due to its rather linear structure, to forming complexes whereas when cytoplasmic fraction is prepared, the predominant species is rRNA. This observation suggests a general method for the isolation of or the enrichment in this type of RNA from any cells or tissues where it is expressed: TRIzol extraction followed by isolation of DNA fraction and subsequent DNase treatment.

The enzymatic activity central to the mRNA amplification process is RdRp; in mammalian cells, it was first detected in anemic rabbit reticulocytes (Downey et al., 1973). It is now apparent that it operates in most, if not all, mammalian cells (Ahlquist, 2002; Iyer et al., 2003; Kapranov et al., 2010), however, the outcomes are qualitatively different in cells undergoing erythroid differentiation, where full-length antisense transcripts of mRNA are produced (Volloch et al., 1996), and in other studied cell types where only short antisense transcripts are generated (Iyer et al., 2003; Kapranov et al., 2010). It is possible that the “omnipresent” RdRp activity is the core enzyme that lacks a processivity component. It follows that in special circumstances, such as erythroid differentiation in particular and possibly terminal differentiation in general, or in some pathologies, what is induced is not, or mainly not, the core enzyme, but rather its processivity factor.

It appears that mRNA amplification is specific for certain physiological situations and, in the case of erythroid differentiation, even for specific stages of differentiation. In our experiments, most of the spleen cells at the time of harvesting were basophilic erythroblasts, a stage when hemoglobin rapidly accumulates and its levels reach their peak (Wojda et al., 2002), as do levels of pepRNA. Subsequently, hemoglobin levels are maintained at a plateau through the reticulocyte phase (Wojda et al., 2002) whereas the levels of pepRNA drastically decline. Therefore, it appears that globin mRNA amplification is active during the hemoglobin accumulation stages and becomes redundant during the maintenance period.

The detection of non-modified chimeric sequences, together with previous observation of non-modified antisense molecule (Volloch et al., 1996), suggests that nucleotide modifications are introduced posttranscriptionally and occur only within the sense component of a double-stranded intermediate. Considering that the only general feature strictly specific for the sense strand is its 3’poly(A) tail, it can be suggested that the poly(A) region of the sense component of double-stranded RNA is both a recognition site and a starting point for a separating/modifying activity. Assuming that modifications facilitate strand separation, taking into account the presence of the terminal 3’poly(A)/5’poly(U) sense/antisense segment in a double-stranded complex and considering the necessity to prevent its snapback after strand separation, it is likely that one of the modified nucleotides is an altered adenosine. This possibility is consistent with the slow rate of pCp ligation and with the observation that the modified pepRNA species does not bind to oligo(dT), a feature offering a simple method for its separation from conventional mRNA species. Moreover, an independent argument can be made that the two modified nucleotides are altered adenosine and guanosine. Only these two bases have the same mass number differential, 16, as do the two modified bases, consistent with the possibility that the same modifying group with a mass of 37 is appended to both A and G. The fact that both are purines indicates that the modification likely occurs at a position(s) not available or not present in pyrimidine nucleotides. One likely candidate is purine position 3 in proximity to the Watson-Crick interphase, whose equivalent is used in pyrimidines for sugar attachment. It should be emphasized that while the delineation of the chemical nature of nucleotide modification(s) remains important, accomplishing this will not, on its own, augment our understanding of the origin and the identity of the RNA of interest, but will pave the way to removing or manipulating these modifications so as to allow direct sequence analysis.

Considering, as discussed above, that strand separation/nucleotide modification commences at the 3’poly(A) of the sense strand and proceeds in the 5’ direction, it can be suggested that when the helicase complex reaches a single-stranded stretch at the head of a pinhead structure it cleaves at or close to its 3’ end leaving the 5’end with (OH) and the 3’ end with a phosphate group. This is consistent with the detection of a class of non-modified antisense globin RNAs lacking their 3’-terminal complementary element, i.e. presumably cleaved at the 3’ end of single stranded loop (Volloch et al., 1996). One of the consequences of such a mechanism is that since the cleavage is the ultimate act in the generation of the chimeric end-product of RNA amplification, it is formed *already* modified and, unlike its pinhead chimeric precursor, is *never* present in the unmodified form and thus cannot be detected by sequencing methods dependent on the lack of modifications.

What is special about globin RNA that makes it the major, if not the only, amplification target in erythroid cells? Whereas the specificity of genome-based transcription is centered on the 5’ end of a transcript, this seems to be not the case for mRNA amplification. Our earlier study (Volloch et al., 1996) showed that antisense synthesis initiates within the 3’ poly(A) of mRNA molecule and suggested that any 3’ poly(A)-containing RNA might serve as a template for an antisense strand synthesis by RdRp (Volloch et al., 1996). Observations by Kapranov and co-workers, who showed a widespread synthesis of antisense RNA initiating apparently indiscriminately at the 3’ poly(A) of mRNAs in human cells (Kapranov et al., 2010), expanded and generalized this notion. The RNase H mapping analysis described here indicated that pepRNA terminates at the 3’end with a truncated poly(A), reflecting short 5’poly(U) of the antisense strand (Volloch et al., 1996). Taken together, the above observations suggest that RdRp can initiate within 3’poly(A) of any or most RNA molecules. Specificity of the amplification process appears to be determined at the 3’end of the antisense strand by its ability to appropriately fold and self-prime synthesis of a sense strand.

In the course of our analysis we identified sense globin RNA sequences with non-conventionally templated 5’-terminal Us added to 5’UTR truncated at positions corresponding to potential cleavage sites within the antisense component of the chimeric intermediate. What could be the origin of such sequences? Provided the presence of 3’poly(A) on an RNA molecule is necessary and sufficient for initiation of RNA-dependent RNA synthesis, another mRNA amplification paradigm may be considered. If an antisense transcript is polyadenylated at the 3’ end by a known or a novel cytoplasmic poly(A) polymerase, it would become a valid template for RdRp. Since the antisense strand has, by virtue of initiation within the poly(A) of conventional mRNA, a poly(U) stretch at the 5’ end, its transcription by RdRp would result in a sense strand with poly(U) at the 5’ end and poly(A) at the 3’ end, also a legitimate RdRp template. Since strand separation mechanisms are in place and the described sequence of events can occur repeatedly, the process will amount to an intracellular polymerase chain reaction, iPCR. The obvious question regarding such a process is its specificity. If 3’ polyadenylation of an antisense molecule is coupled with cleavage of the chimeric pinhead intermediate, the specificity of iPCR would be equal to that of the initial chimeric amplification round. Since both strands would posses 3’poly(A), strand separation and associated nucleotide modification would start at both ends of a linear double stranded structure and detection of unmodified 5’poly(U)-containing sense strand would be more likely than that of unmodified 3’poly(A)-containing antisense strand. If iPCR indeed operates in conjunction with the chimeric pathway, such a two-tier mRNA amplification mechanism may lead to interesting “asymmetrical” results, for example CTF of a protein produced in the chimeric round and the same complete protein generated through the iPCR pathway. Moreover, during the chimeric round, antisense self-priming may occur in such positions as to result in translationally non-functional sense strand fragments. However, regardless of the outcome of the chimeric round, the resulting 3’-truncated/polyadenylated antisense strand could give rise, via iPCR process, to a 5’-truncated/polyuridylated sense strand with complete mRNA coding content. In fact, such a mechanism could explain the observation by Kapranov and co-workers (Kapranov et al., 2010) of a class of 5’-polyuridylated mRNA molecules typically lacking 14-18 genome-encoded 5’-terminal nucleotides. In terms of the mRNA amplification/iPCR model, these truncations reflect the average size of the 3’-terminal complementary/priming element of an antisense strand.

The potential physiological significance of the mRNA amplification, with every genome-originated mRNA molecule acting as a possible template, can hardly be overstated. Malfunctions of this process may be involved in pathologies associated either with the deficiency of a protein normally produced by this mechanism or with the overproduction of a protein normally not involved in such a process. As was mentioned above, if self-priming of the antisense strand occurs within the segment corresponding to the coding region of mRNA, the resulting amplified protein product could be a C-terminal fragment of the original protein, a mechanism proposed to underlie precursor-independent generation of beta-amyloid in sporadic Alzheimer’s disease (Volloch, 1996; 1997).

In the framework of the RNA amplification model, the best conceivable experiment to assess its physiological significance would be to interfere *in vivo* with the extent of complementarity between the two elements involved in self-priming. In fact, multiple experiments of this sort have been carried out by nature. Thus, at least four different types of familial thalassemia (of the mild beta+ type, characterized by reduced production of beta chains) are associated with different point mutations in the 5’UTR of human beta globin mRNA that impede the amplification-associated complementary relationship described above and appear to solely account for the disease (reviewed in Cao and Galanello, 2010).

## METHODS

Balb/C mice, 2-4 month old, were rendered anemic by daily intraperitoneal injections of 0.1 ml of 0.8% neutralized phenylhydrazine as described (Bastos et al., 1977). After 6 days, 80-90% of the cells in circulating peripheral blood were reticulocytes and a similar proportion of spleen cells were erythroid, most having the characteristics of basophilic erythroblasts, consistent with previous observations (Bastos et al., 1977; Conkie et al., 1975). Spleen and blood were collected on day seven as described (Aviv et al., 1975). For RNA preparation minced spleen was passed sequentially through sieves of mesh 100, 150 and 200 or through 100 μm cell strainer (BD Falcon). Cells were collected by centrifugation and lysed in a buffer containing 30 mM Hepes, pH8.2; 50 mM NacCl; 5 mM MgCL_2_; 1% NP40; 10% sucrose; 5 mM DTT and 0.5 units/μl RNase inhibitor (Promega, NEB). After centrifugation SDS and EDTA were added to 0.2% and 10 mM, respectively, and the lysate was extracted with phenol/chloroform with or without prior treatment with proteinase K. Precipitated RNA was dissolved, treated with DNase, phenol extracted and re-precipitated. Reticulocytes were lysed by incubation and occasional pipetting in 3 mM MgCl_2_; lysis was indicated by a change of coloration, at which point NaCl was added to 100 mM. After centrigugation the lysate was adjusted to 100 mM Tris/HCl pH 7.5; 50 mM NaCl; 0.2% SDS and 10 mM EDTA and extracted as described above. Reticulocyte globin RNA was purified by oligo(dT) chromatography followed by high stringency hybridization with a mixture of immobilized α and β globin-specific probes (probes α3, α7, α10, β3, β7 and β10 listed below). For immobilization and use of oligonucleotides, an oxidizable solid support (Molecular Biosystems, San Diego) was employed following manufacturer’s instructions (Volloch et al., 1990; 1991).

RNA was resolved by electrophoresis on agarose-methylmercury (Auzabel et al., 1987; Sambrook et al., 1989) and the major RNA components were visualized by staining with ethidium bromide. The RNA of interest was electroeluted from slices of gel in electrophoresis running buffer in dialysis bags (Spectrapore CE, MWCO 3500). Slices of agarose-methylmercury gels were soaked in 0.1 M DTT for 30 min prior to electroelution (Lemishka et al., 1981). Eluates were passed trough Spin-X centrifugation units (Costar) and dialyzed against 1 mM EDTA, pH 6.2.

To analyze nucleotide composition, 20 μg of gel-eluted pepRNA and of poly(A)-containing RNA eluted from a slice of the same gel located between rRNA and pepRNA bands (control RNA) were dissolved in 20 μl of water, denatured by 3 min incubation at 100 ˚C, adjusted to 10 mM ammonium acetate pH 5.3 and treated with nuclease P1 (EC 3.1.30.1; Sigma) for 2 hrs at 45 ˚C (Crain, 1990). Following P1 treatment, the reaction was adjusted to 0.1 M ammonium bicarbonate pH 7.9 and incubated with 2 milliunits of snake venom phosphodiesterase (EC 3.1.4.1; Sigma) at 37 ˚C for 2 hrs (Crain, 1990). Finally, 0.5 units of bacterial alkaline phosphatase (EC 3.1.3.1; Sigma) was added and incubation continued for 1 hr at 37 ˚C (Crain, 1990). The resulting digest was analyzed by reverse phase HPLC on a 2.1 × 250 mm C18 column (Vydec) with a 2.1 × 150 mm silica gel guard column at a flow rate of 1.5 ml/min. Gradient conditions were as described (Pomerantz and McCloskey, 1990). Further analysis of the eluted fractions was carried out on ABI Voyager MALDI-tof mass spectrometer using 2,5-dihydroxybenzoic acid as the matrix. HPLC and mass-spectrometric analyses were performed by Analytical Biotechnology Services, Boston.

For oligonucleotide-mediated RNase H cleavage (RNase H mapping), pepRNA and reticulocyte globin RNA were labeled at either 5’ or 3’ termini. 5’ labeling was carried out with polynucleotide kinase and γ^32^P ATP. Prior to phosphorylation, reticulocyte globin RNA was treated with tobacco acid pyrophosphatase to de-cap RNA molecules and with calf intestinal phosphatase to remove the remaining phosphate. No preliminary treatment was required for 5’-labeling of pepRNA; labeling reactions with both RNA preparations proceeded with similar efficiency. 3’-labeling was carried out by ligation with ^32^P-labeled pCp and, unlike 5’ labeling, proceeded much slower with pepRNA than with reticulocyte globin RNA or with gel-eluted conventional globin RNA from anemic spleen. Labeled RNA was treated with proteinase K, phenol extracted, precipitated and subjected to electrophoresis on 8% polyacrylamide-urea gels. RNA was visualized by authoradiography, eluted by overnight incubation in ammonium acetate 0.5M, pH 7.2; SDS 0.1%; EDTA 1 mM; tRNA 20 μg/ml, extracted with phenol/chloroform and precipitated with ethanol. For RNase H cleavage, 10-50 ng aliquots of RNA were mixed with 100-fold excess of oligonucleotide complementary to different segments of either α or β globin RNAand with 2 μg of unrelated carrier RNA, incubated in 1 mM EDTA pH 6.2 for 3 min at 95οC, cooled on ice, adjusted to 40 mM Tris pH 8.0, 7 mM MgCl2, 150 mM NaCl, and prehybridized for 30 min at 37 ˚C. Reactions were started by addition of BSA to 50 μg/ml, DTT to 5 mM, RNase inhibitor to 1 unit/μl and RNase H and were continued for 60 min at 37 ˚C. RNase H was used at 10-30 milliunits per reaction with globin RNA and 100-500 milliunits per reaction with pepRNA. Following cleavage, the RNA was treated with proteinase K, phenol extracted, precipitated and analyzed by electrophoresis on 8% polyacrylamide-urea gels. When cleavage resulted in a short fragment, samples were analyzed directly, without proteinase K treatment and extraction. The following oligonucleotides were used in RNase H mapping (numbers in parenthesis indicate location of oligonucleotide termini on corresponding mRNA):

α1: GAAGAAGGGCATGGCCAGAAGG (506-485); α2: ACGGTACTTGGAGGTCAGCACG (461-440);

α3: GGCATGTACCGCGGGGGTG (407-389); α4: GTGGTGGCTAGCCAAGGTCAC (377-357);

α5: AAGGTCACCAGCAGGCAGTGG (364-344); α6: CCAGAGAGCACCATGGTTTCTTCCT (49-25);

α7: GGCCACCAATCTTCCCCCAG (96-77); α8: AGCTCCATATTCAGCACCATGGCC (116-93);

α9: GGTCTTGGTGGTGGGGAAGCT (161-141); α10: GATCCACACGCAGCTTGTGGG (321-301);

α11: TCGGCAGGGTGGTGGCTAGCC (385-365); β1: AACAACTGACAGATGCTCTCTTGGGAA (565

-539); β2: GGGGGTTTAGTGGTACTTGTGAGCC (504-480); β3: GGCAGCCTGTGCAGCGGG (443-426);

β4: TCCTTGCCAAGGTGGTGGC (419-401); β5: GCTGTCCAAGTGATTCAGGCCATC (297-274);

β6: ATGATGTCTGTTTCTGGGGTTGTGAGT (56-30); β7: GGTGCACCATGATGTCTGTTTCTGG (64-40);

β8: CCTTTCCCCACAGGCAAGAGACAG (109-86); β9: CCAGGGCCTCACCACCAAC (142-124);

β10: GGCTGGCAAAGGTGCCCTTGAGG (322-300).

For cell-free translation, pepRNA and regular globin RNA from anemic spleen were eluted from agarose-methylmercury gels as described above. Regular globin RNA was further purified and its traces were removed from pepRNA by high stringency hybridization of both preparations with a mixture of immobilized α and β globin-specific probes (α3, α7, α10, β3, β7 and β10). Reticulocyte globin RNA, similarly purified, was also used in this assay. The translation was carried out using wheat germ extract (Promega) following the manufacturer’s instructions in the presence of methylmercury hydroxide as described (Payvar and Shimke, 1979). The translation products were analyzed on 15% polyacrylamide-SDS gels (Bio-Rad). Immunoprecipitation of translation products was carried out as described (Harlow and Lane, 1988) using sheep antisera against adult murine globins (courtesy of Dr. Jeffrey Ross, Univ. of Wisconsin).

For Next Generation Sequencing, cytoplasmic RNA from anemic spleen was used for preparation of sequencing libraries. Two library preparation kits were employed: TruSeq RNA/V2 (Illumina) and NEBNext Ultradirectional RNA (NEB); both kits were used following manufacturers’ instructions with minor modifications. Libraries obtained were analyzed on a MiSeq sequencer (Illumina) as directed by manufacturer. The resulting reads were either aligned with references containing sequences of chimeric junctions or blasted against sequences of chimeric junctions and other sequences of interest at the stringency word 7. Following the analysis of reads, fragments of interest were extracted from raw data and analyzed.

## ACKNOWLEDGEMENTS

Authors are grateful to Philipp Kapranov for insightful and stimulating discussions, to Bruce Schweitzer and Paul Leavis for experimental help, to Matt Vivero, Ugur Ayturk and Nam Pho for help with the analysis of sequencing data, to Matthew Warman for making his resources available to the study, to Jeff Ross for gift of antibodies, to Gili Naveh and Garif Yalak for help with figure preparation and to Wei Huang for dissections of mice.

## AUTHOR CONTRIBUTIONS

VV and SR conceived and/or developed all concepts described in the present study. SR and VV designed and carried out, with the help of individuals listed in the acknowledgements section, all experiments described above. VV wrote the manuscript; SR and BO participated in the writing of the manuscript. BO hosted the later stages of the study and participated in discussions and analysis of the results.

